# MateSim: Monte Carlo simulation for the generation of mating tables

**DOI:** 10.1101/239178

**Authors:** A. Carvajal-Rodríguez

## Abstract

In species with sexual reproduction, the mating pattern is a meaningful element for understanding evolutionary and speciation processes. Given a mating pool where individuals can encounter each other randomly, the individual mating preferences would define the mating frequencies in the population. However, in every mating process we can distinguish two different steps. First, the encounter between partners. Second, the actual mating once the encounter has occurred. Yet, we cannot always assume that the observed population patterns accurately reflect the individual’s preferences. In some scenarios the individuals may have difficulties to achieve their preferred matings, such as in monogamous species with low population size, where the mating process is similar to a sampling without replacement. In the latter, the encounter process will introduce some noise that may disconnect the individual preferences from the obtained mating pattern. Actually, the difference between the mating pattern observed in a population and the mating preferences of the individuals have been shown by different modeling scenarios.

Here I present a program that simulates the mating process for both discrete and continuous traits, under different encounter models and individual preferences, including effects as time dependence and aging. The utility of the software is demonstrated by replicating and extending, a recent study that showed how patterns of positive assortative mating, or marriage in human societies, may arise from non-assortative individual preferences. The previous result is confirmed and is shown to be caused by the marriage among the “ugliest” and oldest individuals, who after many attempts were finally able to mate among themselves. In fact, I show that the assortative pattern vanishes if an aging process prevents these individuals from mating altogether. The software MateSim is available jointly with the user’s manual, at http://acraaj.webs.uvigo.es/MateSim/matesim.htm

## 1. Introduction

In every mating process we can distinguish two key steps. First, the process of encounter between partners which depends on the phenotypic distribution of the population. Second, once the encounter happens, the mutual individual preferences take action by accepting or rejecting the mate.

Yet, we should distinguish between individual mating preferences and the observed population mating patterns. The population mating pattern is a product of (a) the phenotypic distribution of the population and (b) the individual preferences. Accordingly, if the phenotypic distribution remains constant during the mating season, we could predict the mating pattern given the phenotypic distribution and the individual preferences. We expect the phenotypic distribution to be constant in some scenarios such as polygamous or monogamous species with high population sizes. However, in other scenarios such as monogamous species with low population size, the phenotypic distribution varies during the mating season, and so, the mating process is similar to a sampling without replacement (from the point of view of available partners) which means that the encounter process will introduce some bias that disconnect the individual mating preferences from the obtained mating pattern as predicted given the initial phenotypic distribution (Gimelfarb, 1988a, b). Moreover, the difference between the mating pattern observed in a population and the mating preferences of the individuals, has been evidenced by modeling scenarios that produce positive assortative mating without assortative individual preferences (Burley, 1983; Xie et al., 2015).

Most models assume that encounters are random, although depending on the species this random encounter can be sequential i.e. individual by individual, or in groups, leks, etc. In addition, the species may be polygamous (models with replacement) or monogamous, the latter implying that when a pair is formed these individuals are removed from the mating pool.

Mating in a monogamous species may still resemble a sampling with replacement if the population size is high enough to permit that the mating sample is only a small fraction of the whole population, so that the frequencies of available phenotypes are not altered during the mating process. However, if most of the population is involved in the mating, then the process will resemble a model without replacement. In any case, once an encounter happens, mating depends on the individual preferences between partners. Therefore, it is interesting to explore how different encounter models interact with individual preferences, in order to get a better understanding of how the population mating pattern and the individual mating preference are connected (Burley, 1983).

Simulation is a key tool for exploring mate choice evolution scenarios (Carvajal-Rodriguez and Rolán-Alvarez, 2014; Fernández-Meirama et al., 2017; Rolan-Alvarez et al., 2015; Xie et al., 2015). In the present work, I propose a mating simulation software that allows the generation of mating tables under different kinds of traits, encounter rules and preferences. The program may be useful for exploring the relationship between mating preferences and the observed population pattern under different conditions. It may also be useful for testing different estimators of non-random mating and sexual selection, for example, we may simulate several mating tables under a model without preferences that might serve as a null model for hypothesis testing; or we may simulate tables under positive assortative preferences for comparing the power of different estimators (Carvajal-Rodriguez, 2018).

I demonstrate the software by replicating a recent study that utilized a continuous encounter mating (EM) model to show how patterns of positive assortative mating may arise from non-assortative individual preferences (Xie et al., 2015).

## 2. Software description

### 2.1 General algorithm

MateSim performs Monte Carlo simulation of mating tables under different discrete or continuous (unimodal or bimodal) trait models. The general algorithm is quite simple (Fig. 1).

**Fig 1.**
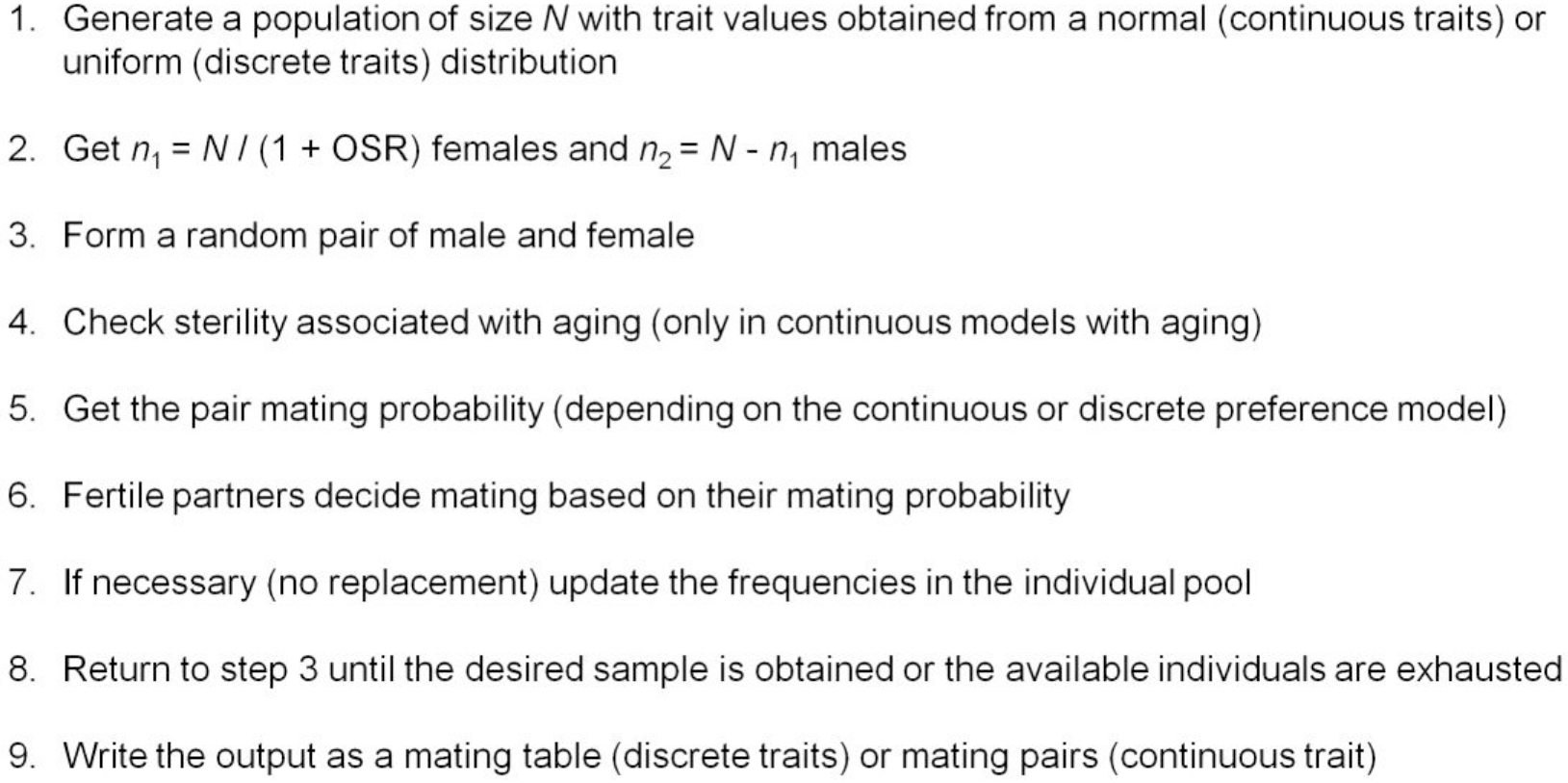
Algorithm for simulating mating pairs. OSR: operative sex ratio.

The first step consists in generating a population of individuals either from a normal distribution (or a mixture of two normal distributions when bimodality is desired) if the mating trait is continuous, or from a uniform distribution, if the mating trait is discrete. For continuous traits, the phenotypic distribution can be bimodal. This is attained by means of a two-component normal mixture (see details in the appendix B).

In addition, the user can also specify any desired phenotypic distribution. In this case, step 1 is skipped and the phenotypic distribution should be read from a file in the correct format (see the program manual). Next, individuals are separated into two sexes as given by the operational sex ratio, which is the ratio of sexually competing males and females that are ready to mate (Clutton-Brock, 2007).

The following step is a key point in the algorithm and refers to the way in which the random encounter occurs between individuals.

### 2.2 Pair formation process

This corresponds to the step 3 in the mating algorithm (Fig. 1). I distinguish two main processes of pair formation: with or without replacement of mating types. When the availability of individuals is not affected by matings that have already occurred, the expected mating pattern in the population is completely determined by the population frequencies and the individual preferences. Thus, the process of pair formation can be viewed as sampling with replacement (SR models) from the set of population types. Polygamous species or monogamous in which only a small fraction of the population mate during a given mating season, are adequately described by this type of pair formation.

On the contrary, when matings are monogamous and most of the population does actually mate, the proportion of available individuals for mating must be updated after each mating event (sampling without replacement from the pool of population types). Processes of the latter kind are the so called encounter-mating (EM) models (Gimelfarb, 1988b).

MateSim simulates mating patterns under both types of pair formation. In the case of the EM models I distinguish individual EM, in which only one encounter takes place at a time, and mass EM, in which more than one pair can be formed simultaneously (Gimelfarb, 1988b). See Tables S1 and S2 in the manual of the program for a detailed explanation on how to define the different pair formation models.

### 2.3 Aging in continuous models

When the preference model is for continuous traits, i.e. the matings are not tabulated by phenotypic classes, we take into consideration each given individual in the population.

In this case, after the pair formation we may evaluate the aging state of the pair. This corresponds to step 4 in Fig. 1. Aging is optional, if the model was defined without aging this step is omitted.

However, it seems realistic to consider that, independently of the mating trait, the mating probability of an individual decreases with age. The latter may occur because the individual becomes old or sterile or simply because he/she does not have enough energy to invest in mating or because the mating season ends.

Without loss of generality, when an individual is not being able of mating because of aging, we say that the individual is sterile. I model the aging process regarding to mating, by means of the Weibull distribution. This distribution is versatile and adequate for modeling survival and time-to-failure processes. In our context, the survival function *R*(*t*) represents the probability that at age *t* an individual is not yet sterile.

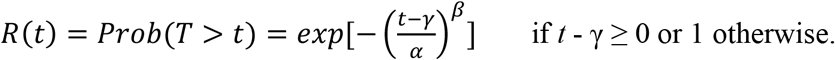

The random variable *T* represents the age to sterility, γ is the age starting from which an individual may be sterile; α represents the 63-percentile, i.e., the age (counted from age γ i.e. α = *t* − γ) at which 63% of the population is sterile, and β is a shape parameter that stands for the increasing risk of being sterile. Thus, if β = 2, the increase of sterility is linear with age while if β > 2 the risk of sterility increases faster at higher age. Age is measured in terms of mating attempts until married (EM models) or until the mating season ends (sampling with replacement model). Thus, I define the age of an individual as the natural logarithm of the mating attempts. Note that each mating attempt requires a new random encounter between any two available individuals of different sex. Therefore, under the EM models, once a pair is established, the age of both partners stops increasing.

After encounter, the aging state of female *i* and male *j* is evaluated. If an individual’s age is above γ then the sterility state is checked prior to the computation of the mating probability. An individual is considered sterile at time *t* when *U* ≥ *R*(*t*), where *U* belongs to *uniform*(0,1). Once an individual is sterile it is discarded from the population. Note that if the model is with replacement, the phenotypic distribution is not affected since it is assumed that the mating individuals are only a small fraction of the population.

### 2.4 Mating under discrete preference models

These models imply that preferences are based on a discrete trait. Preferences are defined as relative mating probabilities (or expected number of matings if not normalized) given an encounter (Fig. 2). The saturated model has *K*-1 parameters where *K* is the product of the number of different female and male types. The saturated individual preference model would produce sexual selection plus assortative mating patterns provided that the individual preferences are reflected in the observed matings (as should occur under SR encounter). Similarly, models with just one parameter may produce female or male sexual selection or positive or negative assortative mating (sexual isolation, Fig. 2). These preference models or any other desired by the user, can be passed as input to the program.

**Fig 2.**
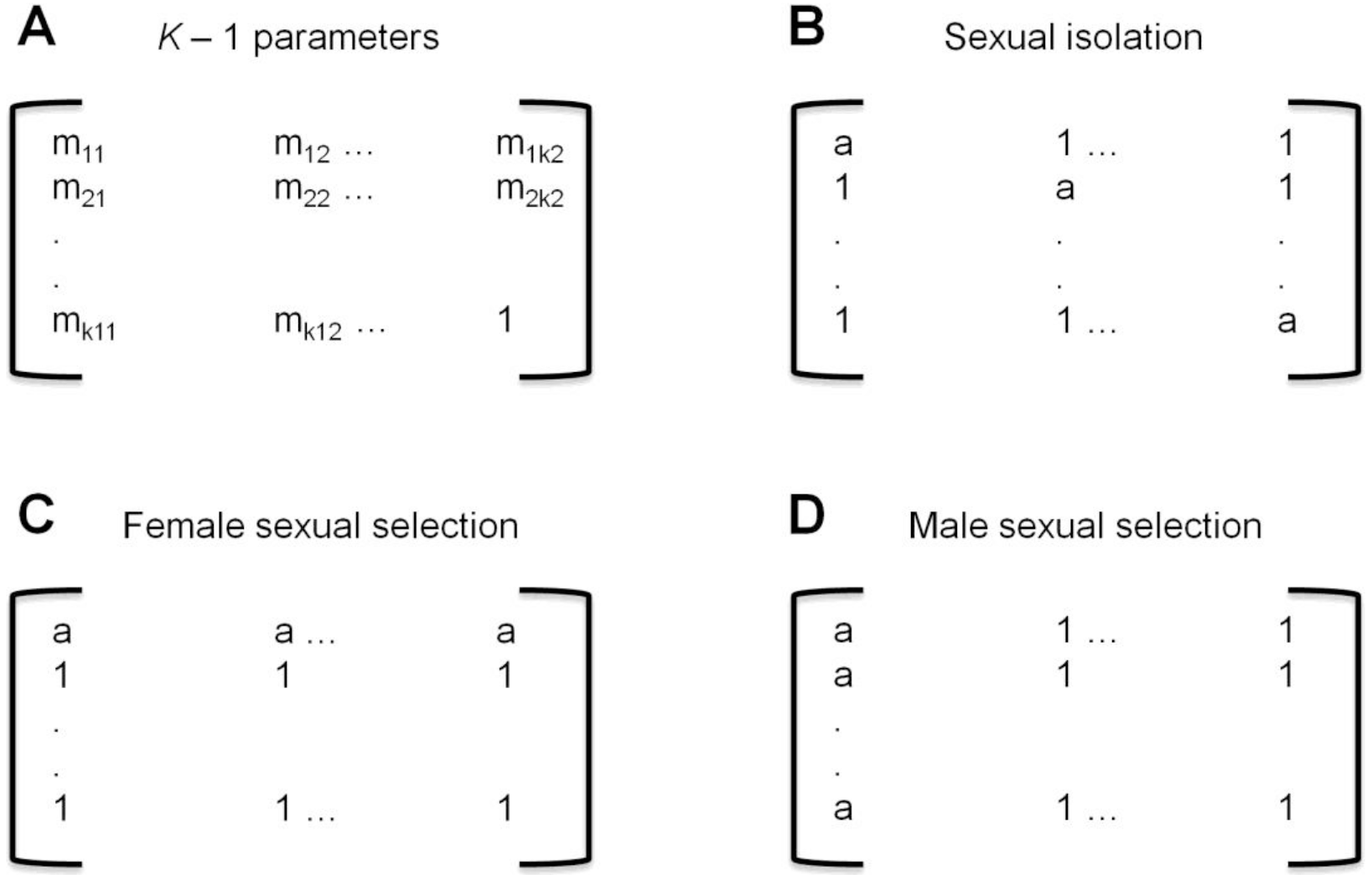
Discrete preference models defined by their expected effects. Females correspond to rows, males to columns.

Simulating matings also requires the population phenotype frequencies, which can be uniform or user defined. Once the population frequencies and the mating preferences are defined, the expected number of occurrences for each mating type *i* × *j* is (Carvajal-Rodríguez, 2018)

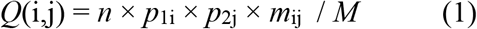

where *p*_1i_ and *p*_2j_ are the *i* female and *j* male population frequencies respectively, *n* is the sample size and *M* = ∑ *p*_1i_ × *p*_2j_ × *m*_ij_. The *m*_ij_ values correspond to the mutual preferences and will depend on the desired model. Note that under EM type models, the phenotype frequencies are updated accordingly after each mating round. The expectation in (1) should be closely met for the SR models, or even for the EM ones when the mating sample size is lower than the given population size. However, when the sample size is close to the population size, the obtained mating pattern can be quite different from the expected.

The format of the obtained mating tables is the same as the JMating (Carvajal-Rodriguez and Rolan-Alvarez, 2006) and InfoMating (Carvajal-Rodríguez, 2018) input files (see the program manual).

### 2.5 Mating under continuous preference models

I implement a general preference function for the continuous model as 
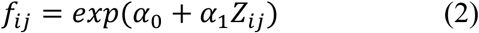

#### 2.5.1 Gaussian models

The Gaussian similarity models (Carvajal-Rodriguez and Rolán-Alvarez, 2014) correspond to the unimodal Gaussian preferences (Lande, 1981) and are obtained by taking α_0_ = 0, α_1_ = −*g*(*C/s*) and Z = (*D*_ij_ − *b*)^2^, where *b* is the bias (see below) and *D*_ij_ is the absolute value of the difference between the female *i* and male *j* phenotypes for the similarity trait. The parameter of the function *g* is a quotient between the choice trait *C* (or its square) and the tolerance *s*, which varies depending on whether the function is FND ((C/s)^2^ Carvajal-Rodriguez and Rolán-Alvarez, 2014) or BD03 ((C^2^/s)^2^ / 2 Bolnick and Doebeli, 2003); the value of *b* is in [0, *D*_max_/2) for positive assortative mating, or fixed to *D*_max_ for negative assortative mating. Given an encounter and the preference function value *f*_ij_, the mating probability for a pair *i*, *j* will be

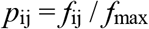

where *f*_max_ is the maximum preference over the available matings. The implementation of the Gaussian preference was performed using a roulette-wheel selection by stochastic acceptance algorithm (Lipowski and Lipowska, 2012).

#### 2.5.2 Logistic models with mutual mate choice

Logistic preference models (Xie et al., 2015) represents open-ended preference functions (Lande, 1981) and may be obtained from (2) by considering the probability for a female *F* (independent of her phenotype) to accept male *j*

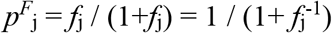

where 
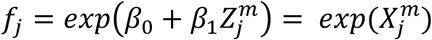

and *Z*^m^_j_ is the trait value of male *j* and *X*^m^_j_ = β_0_ + β_1_*Z*^m^_j_.

Similarly, the probability for a male *M* to accept female *i* is given by

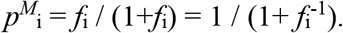

where 
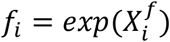

and *X*^f^_i_ = α_0_ + α_1_*Z*^f^_i_. The current implementation of MateSim considers only equal female and male parameters so that α_0_ = β_0_ and α_1_ = β_1_. The mating probability after encounter for a pair *i,j* will be 
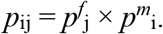

#### 2.5.3 Choice decreasing in time (impatience for mating)

Under both the Gaussian and logistic models above, the choice trait is constant through time. However, we could consider a scenario where individuals become impatient as time passes and an increasing number of matings or marriages are performed, especially if neither polygamy nor divorce is allowed (Xie et al., 2015). Therefore, as time progresses the individuals become less choosy proportional to the number of marriages already performed (see details of the implementation in the appendix A).

## 3. Applications

In the following, I demonstrate the use of the simulator for continuous preference models. I replicate a recent study that utilized a continuous EM model to show how patterns of positive assortative mating, or marriage in human societies, may arise from non-assortative individual preferences (Gale and Shapley, 1962; Xie et al., 2015). I confirm the previous result and demonstrate that the positive correlation is provoked by the marriage among the least-preferred individuals, who after many attempts end up mating among themselves. I furthermore show that, the assortative pattern vanishes in the presence of an aging process.

I use the same parameter values as in the original study. The population consists of *n*_1_ = 5,000 females and *n*_2_ = 5,000 males, and the logistic parameters are α_0_ = −1 and α_1_ = 2. When appropriate, the factor for impatience is c = 0.0005 and finally, I extend the original model by setting the aging process parameters to α = 5, β = 2 and γ = 1. The only difference between this implementation and the original one, is that I allow a maximum of *n*_1_ × *n*_2_ = 25,000 mating attempts. This means that if matings were uniformly distributed, each individual may attempt to mate “only” 5,000 times, which in average, gives a maximum of one attempt with every individual from the other sex. Since the matings are not uniformly distributed and some individuals may spend only a few tries, this means that others may have more than 5,000 attempts. The results are shown in Table 1. Since in the original study all correlation measures gave a similar result, I only utilized one of such measures, namely, the overall correlation over the mating pairs. The model i-EM corresponds to the encounter mating model while the impatient i-EM corresponds to the version with increased probability of mating as time increases. We observe an overall correlation of 0.55 ± 0.001 and 0.44 ± 0.001, which is similar to the values of 0.53 and 0.44 obtained for the same scenarios in the aforementioned study.

I have also considered an encounter mating model in which several pairs can be formed simultaneously (mass-EM), the result is similar (0.42) but the correlation clearly declines (0.16) when the mass model is extended to the impatient version.

**TABLE 1.** Effect of pair formation and aging under a logistic preference function. The population size is 10,000 with equal proportion of males and females. The mating effort refers to the % of the population that is required to mate. The mating success refers to the final number of successful matings.

| Model | Mating<br>effort | Mating<br>success | Correlation | Mean population age<br>(mating attempts) |
| --- | --- | --- | --- | --- |
| SR | 100% | 5000 | $0.001 \pm 0.001$ | 8 |
| i-EM | 100% | 4997 | $0.5458 \pm 0.001$ | 4839 |
| impatient i-EM | 100% | 5000 | $0.4421 \pm 0.001$ | 52 |
| mass-EM | 100% | 5000 | $0.4208 \pm 0.001$ | 209 |
| impatient mass-EM | 100% | 5000 | $0.1553 \pm 0.001$ | 6 |
| i-EM with aging | 100% | 2696 | $0.0548 \pm 0.002$ | 5 |
| i-EM | 90% | 4500 | $0.3740 \pm 0.001$ | 129 |
| i-EM | 80% | 4000 | $0.2502 \pm 0.002$ | 52 |
| i-EM | 60% | 3000 | $0.0775 \pm 0.002$ | 20 |
| i-EM | 50% | 2500 | $0.0377 \pm 0.002$ | 14 |

SR: sampling with replacement; i-EM: individual-encounter mating; mass-EM: mass-encounter mating.

As already commented, it seems more realistic to incorporate an aging component into the model. This is clearly justified in the present case where, when expressing age in terms of mating rounds or attempts, we appreciate that the average age in the i-EM model is almost 5,000 attempts per individual (Table 1). The parameters for the aging model were α = 5, which means that after *e*^5^ ≈ 148 mating attempts, 63% of the individuals are sterile, β = 2, i.e. a linear increase of sterility with age, and γ = 1, which implies that the possibility of sterility begins after 3 attempts. When this aging model is incorporated into the i-EM model, the correlation disappears (0.05), corroborating that assortativeness was caused by an increased homogeneity in the oldest unmarried pool (Xie et al., 2015). The average age, in terms of mating attempts, collapses to 5 which is far less than the 4,839 attempts in the model without aging.

Even if we use a delayed aging model in which the occurrence of sterility is only possible after 55 mating attempts (γ = 4), the obtained correlation is 0.24, which is still far below the value observed without aging. This seems to confirm the pattern, already observed under the model with mating impatience, that the incorporation of more realistic effects, diminishes the observed assortativeness.

Finally, I compare other i-EM models in which not the entire adult population is required to marry. A mating effort of 90% implies that 10% of the adult population remains unmarried. Thus, with 10 or 20% of unmated individuals the correlation falls to 0.37 or 0.25, respectively, and it essentially vanishes when the unmated individuals compress 40-50% of the population (0.08-0.04).

## 4. Conclusions

MateSim is a flexible and versatile tool for simulating different mating processes. It distinguishes between the encounter and mating, and can handle continuous or discrete models. It may be useful for evolutionary studies of mate choice. As a demonstration of the program, I have revisited a study about dynamic processes of marriage in closed systems, and show that after the addition of aging to the original model, the assortative mating pattern obtained without an assortative preference, tends to disappear.

## Acknowledgement

I would like to thank Carlos Canchaya and three anonymous reviewers for their valuable comments on the manuscript. This work was supported by Xunta de Galicia (Grupo de Referencia Competitiva, ED431C2016-037), Ministerio de Economía y Competitividad (CGL2016-75482-P) and by Fondos FEDER (“Unha maneira de facer Europa”).

## Appendix A. Preference functions for continuous traits

### Choice decreasing in time (impatience for mating)

For the Gaussian preference function, “impatience” is implemented as *g*(*C*/ *s*_t_) where *s*_t_ = *s* × (*n*_t_ + 1) is the tolerance *s* multiplied by the number of existing marriages *n*_t_ at time *t* plus one. Thus, the preference (*exp*(−*g*(*C*/ *s*_t_)) tends to 1 independently of *C* as the number of marriages increases. In the current implementation *s*_t_ = *s* by default, this can be changed by means of the tag -*impatience* which if different from 0 defines the impatient behavior (see the program manual).

The impatience for the logistic model is (Xie et al., 2015)

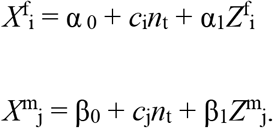

where *c*_i_, *c*_j_ are constants and *n*_t_ stands for the number of existing marriages at time *t*.

In the current implementation *c*_i_ = *c*_j_ = *c* = 0 by default. The value of *c* can be changed, say to 0.001, by the tag

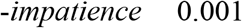

Note that (provided that *c* is positive for the logistic case) as *n*_t_ increases the mating probabilities tend to 1 independently of the mating trait, so mating becomes less selective.

### Aging: Weibull model

The survival function *R*(*t*) represents the probability that at age *t* an individual is not yet sterile.

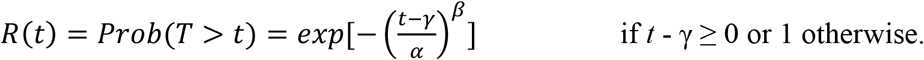

The random variable *T* represents the time to sterility, γ is the age from which on an individual may be sterile; α represents the 63-percentile, i.e. the age (counted from age γ i.e. α = *t* − γ) at which 63% of the population is sterile, and β is the parameter for the increasing risk of being sterile. Then, if β = 2 the increase of sterility is linear with age while if β > 2 the risk of sterility increases faster at higher age. Age is measured in terms of mating attempts until married (EM models) or until the whole mating season ends (sampling with replacement model). Because there can be many attempts, we use a logarithmic scale so that the age of an individual is the natural logarithm of its number of mating attempts.

The implemented continuous preference model by default assumes no age structure, so that *R*(*t*) = 1 for every age. The aging settings can be defined by the tag

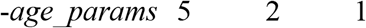

which introduces the values α = 5, β = 2 and γ = 1.

If we do not want the aging process we can simple put a negative sign to the gamma parameter so that

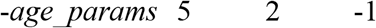

implies that there is not aging (*R*(*t*) = 1 for every age).

An individual is considered sterile at time *t* when *U* ≥ *R*(*t*) where *U* follows an *uniform*(0,1). If the individual happens to be sterile then it is discarded from the population. The iteration ends when the specified or available number of non-sterile individuals have mated.

## Appendix B. Bimodal populations for continuous traits

Consider the density of a two-component normal mixture where *π* is the mixing proportion: *h*(x;μ_1_, μ_2_, σ_1_, σ_2_, *π*) = *π*φ(x; μ_1_, σ_1_) + (1 − *π*)φ(x; μ_2_, σ_2_).

The mixed distribution either bimodal or unimodal, has the mode(s) lying between μ_1_ and μ_2_. The distribution is unimodal or bimodal depending on the values of the three parameters *π*, μ = (μ_2_ − μ_1_) / σ_1_ and σ = σ_2_ / σ_1_ (Robertson and Fryer, 1969). For example, if *π* = 0.5 and σ_2_ = σ_1_ = *s* then *h* is unimodal only if |μ_2_−μ_1_|≤2*s*. More generally, a mixture of two normal densities will not be bimodal unless there is a very large difference between their means, typically larger than the sum of their standard deviations (Schilling et al., 2002). Besides, the number of modes that can be obtained by mixing two normal unidimensional components is 2 and in general for a *D*-dimensional mixture the maximum number of modes is *D*+1 (Ray and Ren, 2012). The modal features of a two component normal mixture remains unchanged under scaling and rotation, so we can generate a bimodal distribution by mixing a standard normal and a normal *N*(μ_2_, σ_2_). In that case, μ_1_= 0, σ_1_ =1, and the condition for bimodality implies μ_2_ > σ_2_ +1.

Therefore, if a continuous bimodal population is desired then the user may define the mixture proportion *π* (0 < *π* <1) and the mean μ_2_ and standard deviation σ_2_ with the restriction μ_2_ > σ_2_ +1 (see command line section in the program manual).

